# LKB1 suppresses growth and promotes the internalization of EGFR through the PIKFYVE lipid kinase

**DOI:** 10.1101/2023.10.19.563158

**Authors:** John R. Ferrarone, Jerin Thomas, Arun M. Unni, Yuxiang Zheng, Michal J. Nagiec, Eric E. Gardner, Oksana Mashadova, Kate Li, Nikos Koundouros, Antonino Montalbano, Meer Mustafa, Lewis C. Cantley, John Blenis, Neville E. Sanjana, Harold Varmus

## Abstract

The tumor suppressor LKB1 is a serine/threonine protein kinase that is frequently mutated in human lung adenocarcinoma (LUAD). LKB1 regulates a complex signaling network that is known to control cell polarity and metabolism; however, the pathways that mediate the tumor suppressive activity of LKB1 are incompletely defined. To identify mechanisms of LKB1- mediated growth suppression we developed a spheroid-based cell culture assay to study LKB1- dependent growth. Using this assay, along with genome-wide CRISPR screens and validation with orthogonal methods, we discovered that LKB1 suppresses growth, in part, by activating the PIKFYVE lipid kinase, which promotes the internalization of wild-type EGFR. Our findings reveal a new mechanism of regulation of EGFR, which may have implications for the treatment of *LKB1*-mutant LUAD.

## INTRODUCTION

The tumor suppressor gene *LKB1* (*STK11*) encodes a serine/threonine protein kinase that is inactivated in several types of human cancers, including up to 30% of lung adenocarcinomas (LUAD) (Shackelford and Shaw, 2009). Tumors with inactivating mutations in *LKB1*, or genomic loss of *LKB1*, are less responsive to both chemo- and immuno-therapy, compared to tumors with wild-type *LKB1* (Krishnamurthy et al., 2021). It is unclear how alterations in *LKB1* promote tumorigenesis. Two studies using mouse models of lung cancer determined that the salt-inducible kinases (SIKs) are required for LKB1-dependent tumor suppression (Hollstein et al., 2019; Murray et al., 2019). Downstream of the SIKs, the CREB-regulated transcriptional coactivators (CRTCs) may play a role in inhibiting growth (Hollstein et al., 2019); however, it is unclear whether the CRTCs specifically mediate the tumor suppressive effects of LKB1, or if they are essential for tumor growth in general.

To determine the mechanisms of LKB1-mediated growth control, we developed a spheroid-based cell culture assay that recapitulated the growth suppressive effects of LKB1 in animal models. We then conducted genome-wide loss-of-function CRISPR screens in spheroid culture and found that LKB1 opposes growth by activating the PIKFYVE lipid kinase. Further, we have determined that activation of PIKFYVE results in the internalization of wild-type (WT) EGFR.

## RESULTS

### Expression of *LKB1* limits growth of *LKB1*-null LUAD lines in spheroid culture, but not in two-dimensional culture

To interrogate the function of LKB1 as a tumor suppressor, we used a retroviral vector to express WT *LKB1* stably in an *LKB1*-null LUAD line (A549, Figure S1A). As expected, expression of *LKB1* led to a marked reduction in the growth of this line as subcutaneous xenografts in mice, compared to *LKB1*-null control cells containing an empty vector (EV) (Figure 1A). However, the expression of *LKB1* had no effect on the growth of this line in two-dimensional (2D) culture (Figure 1B).

**Figure 1:**
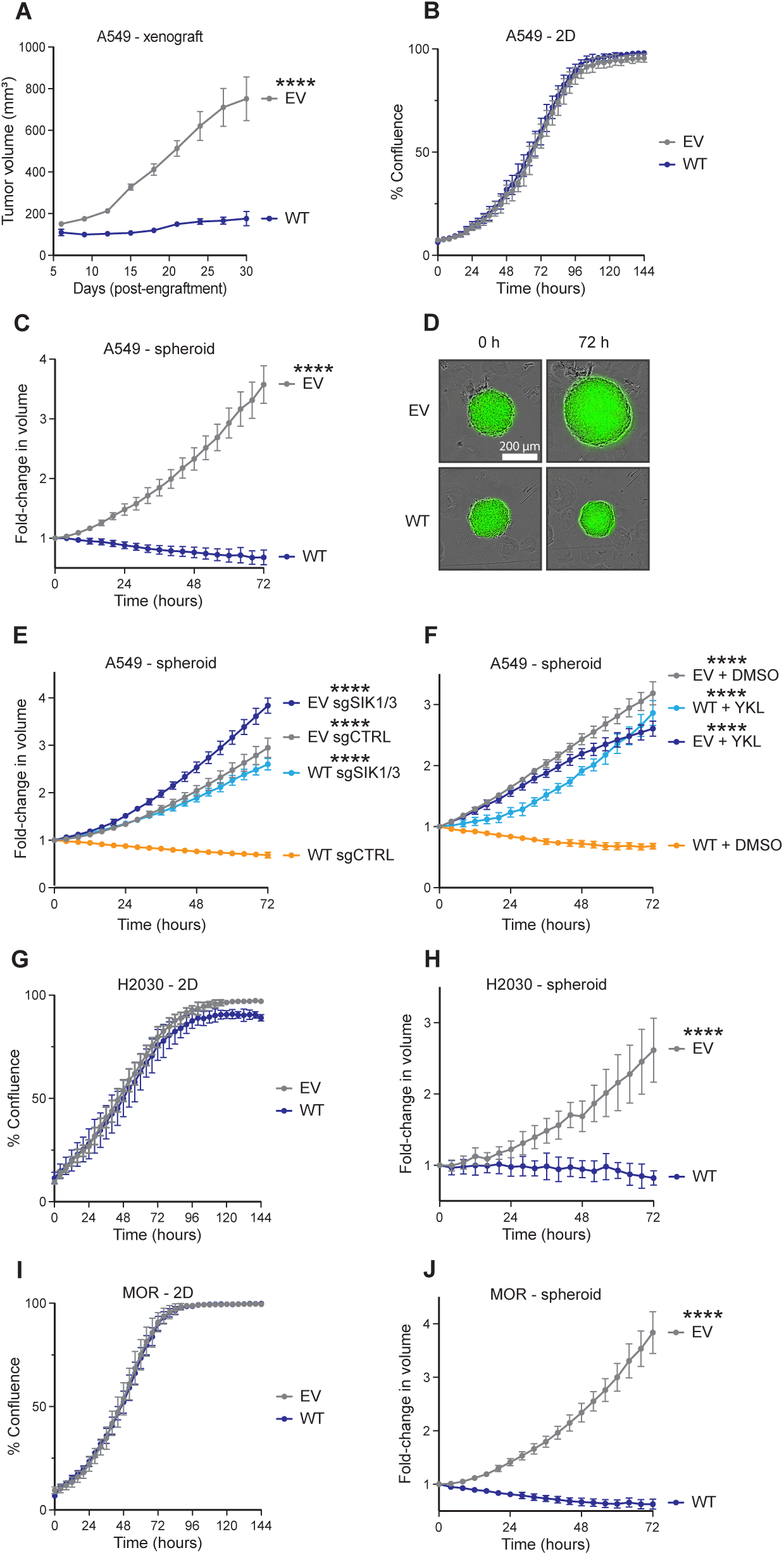
Expression of *LKB1* limits growth of *LKB1*-null human LUAD lines in spheroid culture, but not in two-dimensional culture. **(A)** Growth of A549 cells, which contain an empty vector (EV), leaving the cells *LKB1*-null, or a vector expressing wild-type *LKB1* (WT), as xenografts in the flanks of athymic mice. Tumor volume was measured every 3 days. Error bars indicate standard error of the mean. n = 5. **(B)** Growth of A549 EV and WT cells in 2D culture. **(C, D)** Growth of A549 EV and WT cells in spheroid culture. Growth of the spheroids over 72 hours is shown in panel **(C)** and representative images of the spheroids are shown in panel **(D)**. **(E)** Spheroid growth of A549 EV and WT cells containing a control guide RNA (sgCTRL), or sgRNAs against *SIK1* and *SIK3*. **(F)** Spheroid growth of EV and WT cells treated with DMSO or 2000 nM of the SIK inhibitor YKL-05-099 (YKL). **(G-J)** Growth of H2030 and MOR EV and WT cells in 2D and spheroid culture. Growth measurements were obtained every four hours using an Incucyte S3 live cell imager. For the spheroid culture experiments, growth is expressed as the fold-change in the volume of each spheroid. For the 2D culture experiments, growth is expressed as the percentage of the imaging field that is occupied by cells (i.e., % confluence). Error bars indicate standard deviation. n = 3-6. **** indicates p<0.0001. P-values indicate pairwise statistical comparisons to “WT sgCTRL” or “WT + DMSO” in panels **(E, F)**.

We then evaluated LKB1-dependent growth using a spheroid-based cell culture method. When A549 cells were grown as spheroids in a matrix formed from methylcellulose, re- introduction of *LKB1* dramatically reduced cell proliferation, relative to the *LKB1*-null control cells (Figure 1C, D). We also determined that the kinase activity of LKB1 is required for growth suppression by expressing *LKB1-K78I* (Mehenni et al., 1998), a kinase inactive (KI) version of *LKB1* (Figure S1B), which had no effect on growth in either 2D or spheroid culture (Figures S1C, D). In agreement with the findings in mouse models (Hollstein et al., 2019; Murray et al., 2019), the knockout of *SIK1* and *SIK3* with CRISPR/Cas9, or the treatment of spheroids with YKL-05-099 (Sundberg et al., 2016), a small molecule inhibitor of the SIKs, led to complete restoration of the growth of spheroids with *LKB1*, while having minimal effect on the growth of *LKB1*-null spheroids (Figures 1E, F). We also expressed WT *LKB1* in two additional *LKB1*-null LUAD lines (H2030 and MOR; Figures S1E, F) and confirmed that LKB1 limited the growth of these lines in spheroid culture, but had no effect in 2D culture (Figures 1G-J). These data suggest that spheroid-based cell culture recapitulates the tumor suppressive function of LKB1, as we observed with the subcutaneous xenografts.

We then attempted to recapitulate the tumor suppressive activity of LKB1 in 2D culture by modulating the levels of nutrients in the media. Since the withdrawal of glucose in 2D culture led to activation of LKB1-mediated signaling (as measured by phosphorylation of AMPKα and ULK1) only in cells with WT *LKB1* (Figure S2A), we reasoned that *LKB1*-expressing cells may exhibit reduced proliferation only under low glucose. However, reducing the levels of glucose, amino acids, or serum in the culture media had similar inhibitory effects on the growth of *LKB1*- null and -WT cells in 2D culture (Figures S2B-D). Additionally, we tested the growth of these cells on plates with physiologic levels of surface tension, since mechanical forces can influence the activity of growth-promoting pathways (Vining and Mooney, 2017). However, this also did not lead to a selective reduction in the growth of *LKB1*-WT cells (Figure S2E). Thus, differences in nutrient availability or surface tension do not account for LKB1-dependent discrepancies in growth between 2D and spheroid culture.

### LKB1 suppresses the activity of oncogenic signaling pathways in spheroid culture, but not in 2D culture

To further investigate differences in the activity of LKB1 in 2D and spheroid culture, we evaluated the phospho-proteome of *LKB1*-null and -WT cells in 2D and spheroid culture. In spheroid culture, we noticed a reduction in phospho-peptides associated with the mitogen-activated protein kinase (MAPK) pathway and the mammalian target of rapamycin complex 1 (mTORC1) pathway in the *LKB1*-WT cells, relative to the *LKB1*-null cells (Figure 2A).

**Figure 2:**
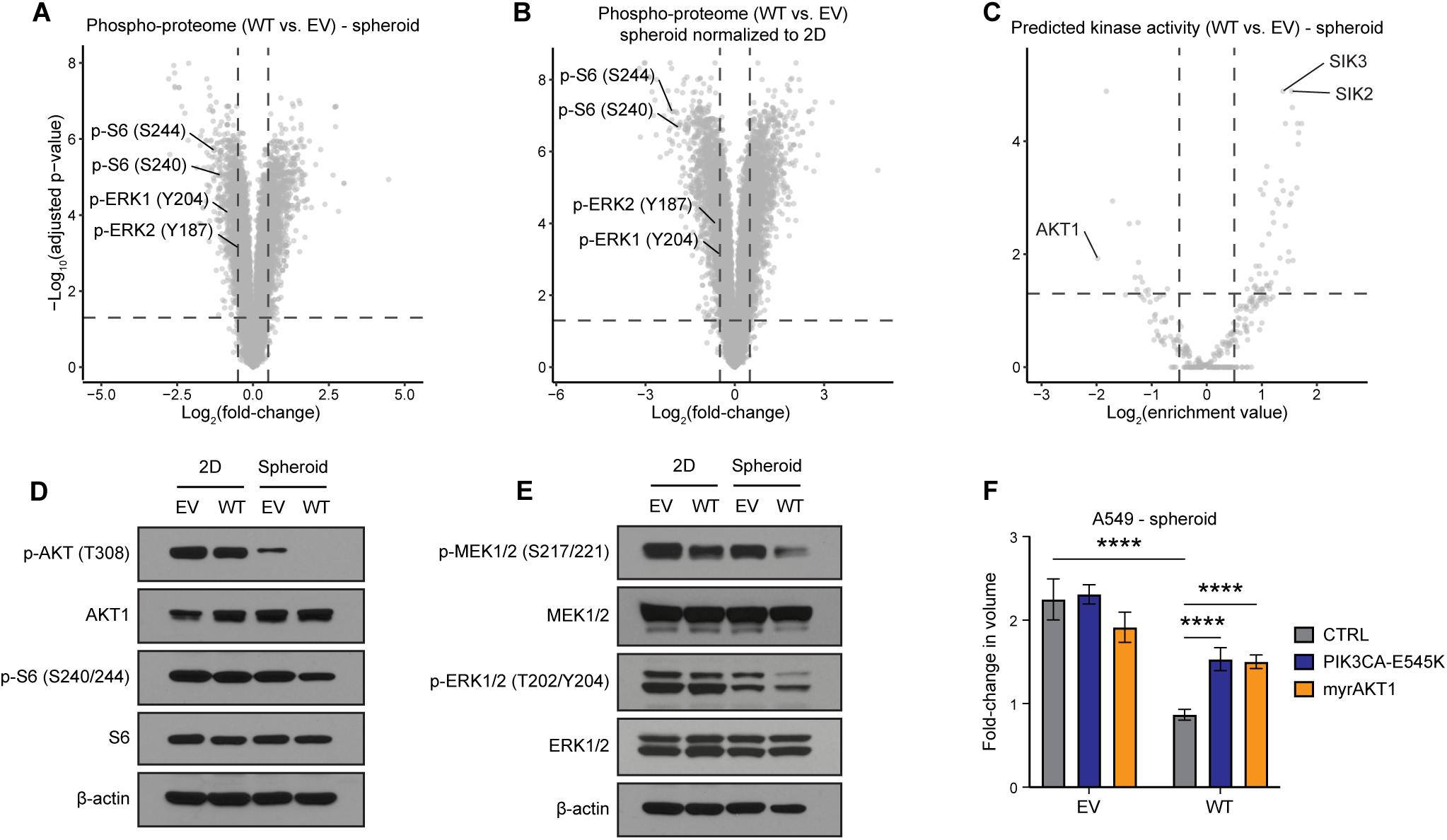
LKB1 suppresses the activity of the mTORC1 and MAPK pathways in spheroid culture, but not 2D culture. **(A)** Relative abundance (expressed as log_2_(fold-change)) of phosphorylated peptides in A549 WT spheroids relative to EV spheroids. **(B)** A comparison of the relative phospho-peptide signal intensities (WT vs. EV) in spheroid culture to the relative values of the same peptides in 2D culture. In **(A, B)**, phospho-peptides that correspond to proteins in the MAPK and mTORC1 pathways are labeled in black. **(C)** The predicted activity of serine and threonine kinases in WT spheroids relative to EV spheroids. **(D, E**) Levels of proteins phosphorylated in the AKT-mTORC1 **(D)** and MAPK **(E)** pathways in EV and WT cells, in 2D and spheroid culture. **(F)** Growth of A549 EV and WT spheroids transduced with an empty lentiviral vector (CTRL) or vectors that express *PIK3CA-E545K* or myristoylated *AKT1* (*myrAKT1*), after 72 hours in culture. Error bars indicate standard deviation. n = 3-6. **** indicates p<0.0001.

To identify changes in signaling that were specific to spheroid culture, we compared the relative phospho-peptide signal intensities (WT vs. EV) in spheroid culture to the relative values of the same peptides in 2D culture. This revealed an even greater reduction in phospho- peptides corresponding to the mTORC1 pathway in the *LKB1*-WT cells relative to the null cells (Figure 2B). We then used a recently developed program that predicts the activity of serine and threonine kinases, using phospho-proteome datasets as input (Johnson et al., 2023). Using normalized phospho-peptide abundance as input (see Figure 2B), we found that the SIKs were among the kinases that were predicted to be most active in cells with LKB1 in spheroid culture, whereas AKT1 kinase activity was predicted to be the most suppressed (Figure 2C). We then confirmed that LKB1 suppresses the activity of the mTORC1 pathway to a greater degree in spheroid culture than in 2D culture (Figure 2D). We also noted that LKB1 suppressed the activity of the MAPK pathway to a greater extent in spheroid culture (Figure 2E).

To assess whether the changes in the AKT1-mTORC1 pathway were functionally significant with respect to growth, we used lentiviral vectors to introduce constitutively active versions of *PIK3CA* (*PIK3CA-E545K*) or *AKT1* (myristoylated *AKT1* (*myrAKT1*)) in *LKB1*-null and -WT cells (see methods). The introduction of *PIK3CA-E545K* or *myrAKT1* partially restored the growth of spheroids with WT *LKB1*, but did not increase the growth rate of *LKB1*-null spheroids (Figure 2F). Together, these results suggest that inhibition of the AKT1-mTORC1 pathway partially accounts for the tumor suppressive activity of LKB1.

### Genome-wide, loss-of-function CRISPR screens in spheroid culture identify genes required for LKB1-mediated tumor suppression

Having established a system to study the tumor-suppressive effects of LKB1, we performed genome-wide CRISPR screens to identify genetic knockouts that enhanced the growth of A549 cells expressing WT *LKB1* (WT) in spheroid culture. As controls, parallel screens were performed in spheroid cultures of *LKB1*-null A549 cells (EV) and in 2D cultures of A549 cells that were *LKB1*-null or that expressed *LKB1*.

To induce genetic knockouts through CRISPR/Cas9, *LKB1*-null or WT cells were transduced with lentivirus carrying the TKOv3 genome-wide CRISPR knockout library (Hart et al., 2017) (Figure 3A). Following selection of infected cells with puromycin, cells were propagated in either 2D or bulk spheroid culture. Genomic DNA was extracted from cells after 21 days in culture and then sequenced to determine the changes in the abundance of DNA sequences encoding single guide RNAs (sgRNAs) over the course of the experiment.

**Figure 3:**
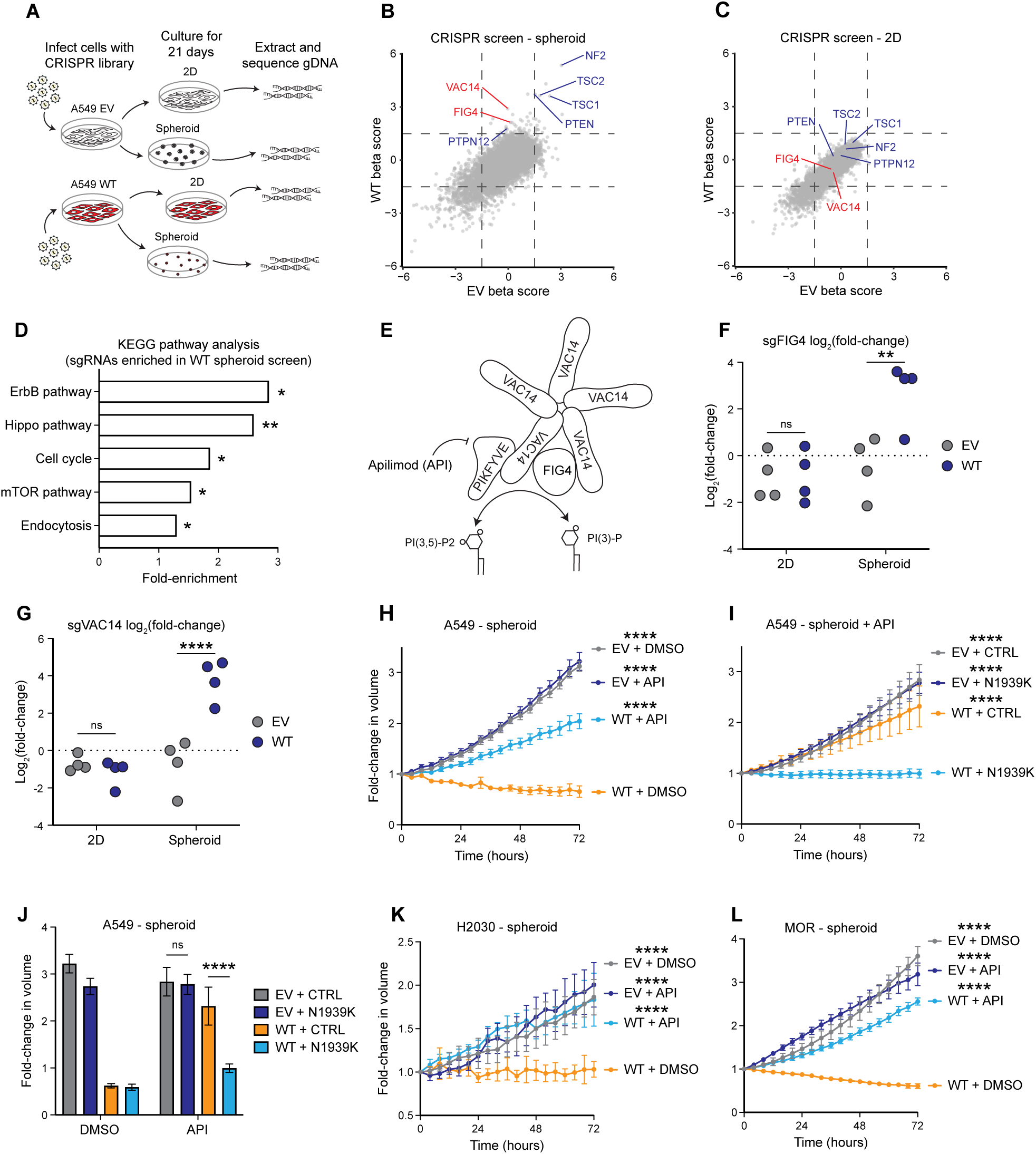
Genome-wide CRISPR screens identify the PIKFYVE lipid kinase complex as a mediator of LKB1-dependent tumor suppression. **(A)** Schematic representation of the CRISPR screen experiment. **(B, C)** Beta score plot of DNAs encoding sgRNAs from cultures of A549 EV and WT cells transduced with a genome-wide CRISPR library and grown in spheroid **(B)** or 2D **(C)** culture. Members of the PIKFYVE lipid kinase complex are labeled in red lettering. Known tumor suppressor genes are labeled in blue lettering. Beta score is a weighted score of the log_2_(fold-change) values of all sgRNAs for a given gene. **(D)** An enrichment analysis of the sgRNAs that were enriched in the WT spheroids (beta score >1) but not in the EV spheroids (beta score < 1). **(E)** A representation of the structure of the PIKFYVE lipid kinase complex (Lees et al., 2020). **(F, G)** Log_2_(fold-change) values of individual sgRNAs corresponding to *FIG4* **(E)** or *VAC14* **(F)** in EV and WT cells grown in 2D or spheroid culture. **(H)** Growth of A549 EV and WT spheroids treated with DMSO or 100 nM of the PIKFYVE inhibitor apilimod (API). **(I, J)** Growth of A549 EV and WT spheroids transduced with an empty lentiviral vector (CTRL) or a vector that expresses a PIKFYVE mutant (N1939K) that is API-resistant, and treated with DMSO or API 100 nM. The growth curve in **(I)** shows the growth of these spheroids in API over a 72-hour period, whereas the bar graph in **(J)** depicts the fold-change in the volume of the spheroids after 72 hours in culture in DMSO or API. **(K, L)** Growth of EV and WT spheroids derived from H2030 **(K)** or MOR **(L)** in the presence of DMSO or API. Error bars indicate standard deviation. n = 3-6. *, **, and **** indicate p<0.05, 0.01, and 0.0001. ns, non-significant. P-values indicate pairwise statistical comparisons to “WT + DMSO” in panels **(H, K, and L)**, and “WT + N1939K” in panel **(I)**.

Strikingly, in the spheroid-based CRISPR screen in cells expressing *LKB1*, we noted a selective enrichment of DNA encoding sgRNAs which correspond to tumor suppressor genes that regulate growth factor receptors (the tyrosine phosphatase *PTPN12*), the Hippo pathway (*NF2*), and the mTORC1 pathway (*PTEN*, *TSC1*, and *TSC2*) (all shown in blue in Figure 3B). Moreover, such enrichment was not observed in cells grown in 2D culture (Figure 3C).

Additionally, a KEGG pathway analysis of the sgRNAs that were selectively enriched in spheroids with WT *LKB1* highlighted the ErbB, Hippo, and mTORC1 pathways (Figure 3D). These findings imply that activation of these pathways may restore the growth of *LKB1*- expressing spheroids and are consistent with a prior study, which showed that LKB1 suppresses growth through the Hippo pathway (Mohseni et al., 2014). These results also suggest that the LKB1-mediated changes in signaling in the MAPK and AKT1-mTORC1 pathways (Figures 2D, E) may be functionally significant.

Of greater interest, we also observed an enrichment of sgRNAs corresponding to two genes involved in the regulation of 5’ phosphoinositides: *VAC14* and *FIG4* (shown in red in Figure 3B). *VAC14* codes for a scaffold protein and *FIG4* encodes a 5’ phosphoinositide phosphatase that form a complex with PIKFYVE – a 5’ phosphoinositide kinase (also known as phosphatidylinositol-3-phosphate 5-kinase type III or PIPKIII) (Figure 3E) (Lees et al., 2020).

Together, this complex regulates the synthesis and turnover of phosphatidylinositol 3,5- bisphosphate (Duex et al., 2006a; Duex et al., 2006b; Jin et al., 2008; Shisheva, 2001), but has not been previously implicated in tumor suppression. The PIKFYVE complex also regulates endosomal trafficking (Posor et al., 2022), which is associated with one of the processes highlighted by the pathway analysis (endocytosis; Figure 3D). Closer inspection of the results of the CRISPR screen showed that most of the individual sgRNAs for these genes were enriched in the *LKB1*-WT spheroids and either depleted or unchanged in spheroids lacking LKB1 (Figures 2F, G). In a replicate of these screens, DNA encoding sgRNAs against *VAC14* was again enriched in *LKB1*-WT cells grown in spheroid culture (Figure S3A).

### LKB1 mediates growth suppression through the PIKFYVE complex

To validate the results of the screen, we knocked out *FIG4* or *VAC14* in the A549 and MOR LUAD lines (Figures S3B-E). We then confirmed that knockout of these genes modestly increased the growth of spheroids with WT *LKB1* (Figures S3F-I) in a three-day assay (as compared to the screen, which was carried out over 21 days). Although DNA encoding sgRNAs against *PIKFYVE* was not enriched in the CRISPR screen of spheroids with *LKB1*-WT, we tested if PIKFYVE also mediates growth suppression, since FIG4 and VAC14 are both known to regulate the activity of PIKFYVE (Lees et al., 2020).

We initially attempted to knock out *PIKFYVE* with CRISPR/Cas9, but we found that none of the four sgRNAs that we tested reduced PIKFYVE protein levels. As an alternative approach, we used the PIKFYVE inhibitor apilimod (Cai et al., 2013) (Figure 3E) to ask whether the enzymatic activity of PIKFYVE affects spheroid growth. We found that apilimod clearly enhanced the growth of spheroids containing WT *LKB1*, but had no effect on the growth of spheroids lacking LKB1 (Figure 3H). To verify that apilimod was acting on its known target, we expressed a mutant version of *PIKFYVE* (which is resistant to the inhibitory effect of apilimod (Gayle et al., 2017)) in the *LKB1*-null and -WT cells. In spheroids with *LKB1*, this PIKFYVE mutant impaired the restoration of growth in the presence of apilimod (Figure 3I). Furthermore, the PIKFYVE mutant had no effect on the growth of the *LKB1*-null spheroids under any condition, which suggests that LKB1 is required for the activation of PIKFYVE (Figure 3J). We then confirmed that apilimod restores the growth of spheroids with WT *LKB1* derived from two other LUAD lines (H2030 and MOR; Figures 3K, L). Additionally, we used apilimod to treat three LUAD lines that harbor endogenous WT *LKB1* (Calu-1, H1792, and SW1573). We found that apilimod augmented the growth of all three lines as spheroids (Figures S4A-C), indicating that PIKFYVE also behaves as a tumor suppressor in cells with physiologic expression of *LKB1*.

To gauge whether LKB1 directly or indirectly mediates the activation of the PIKFYVE complex, we used a program developed by Johnson et al (Johnson et al., 2023) to predict the kinases for all previously reported phospho-serine or -threonine residues on the PIKFYVE complex. Through this approach, we identified numerous sites on PIKFYVE (a 240 kDa protein) that were predicted to be phosphorylation targets of the AMPK-related kinases (Figure S5A) – a family of closely-related enzymes that are directly activated by LKB1, including the SIKs (Shackelford and Shaw, 2009). Closer inspection of these results revealed 6 sites on PIKFYVE that were predicted to be substrates of the SIKs (Figure S5B). One of these sites, Ser307, was previously confirmed to be a target of AMPK (Liu et al., 2013a), and phosphorylation was shown to stimulate the lipid kinase activity of PIKFYVE (Liu et al., 2013a). Given that AMPK and the SIKs are known to share substrates (Shackelford and Shaw, 2009), it is possible that LKB1 may activate PIKFYVE indirectly through the SIKs, as well as through AMPK.

### LKB1 promotes the internalization of EGFR through PIKFYVE

Since PIKFYVE is known to regulate the degradation of growth factor receptors (including EGFR and MET) (de Lartigue et al., 2009; Er et al., 2013), and given that our CRISPR screen highlighted genes involved ErbB signaling as potential mediators of LKB1-dependent tumor suppression, we investigated whether LKB1 influenced the internalization of EGFR through PIKFYVE. To do this, we used a pH-sensitive dye that is conjugated to EGF (pHrodo EGF), which fluoresces when exposed to the acidic environments of the endosomal and lysosomal compartments to indicate that EGF-EGFR complexes have been internalized (Suprynowicz et al., 2010). When we provided *LKB1*-null and -WT spheroids with pHrodo EGF, we noted that fluorescence signal intensity was markedly higher in the *LKB1*-WT spheroids relative to the *LKB1*-null spheroids (Figure 4A, B). Pretreatment of these spheroids with apilimod (a PIKFYVE inhibitor), YKL-05-099 (a SIK inhibitor), or chloroquine (an inhibitor of endocytosis) reduced the fluorescence of *LKB1*-WT spheroids fed with pHrodo EGF (Figure 4C). These results suggest that LKB1 promotes the endocytosis of EGFR through both the SIKs and PIKFYVE.

**Figure 4:**
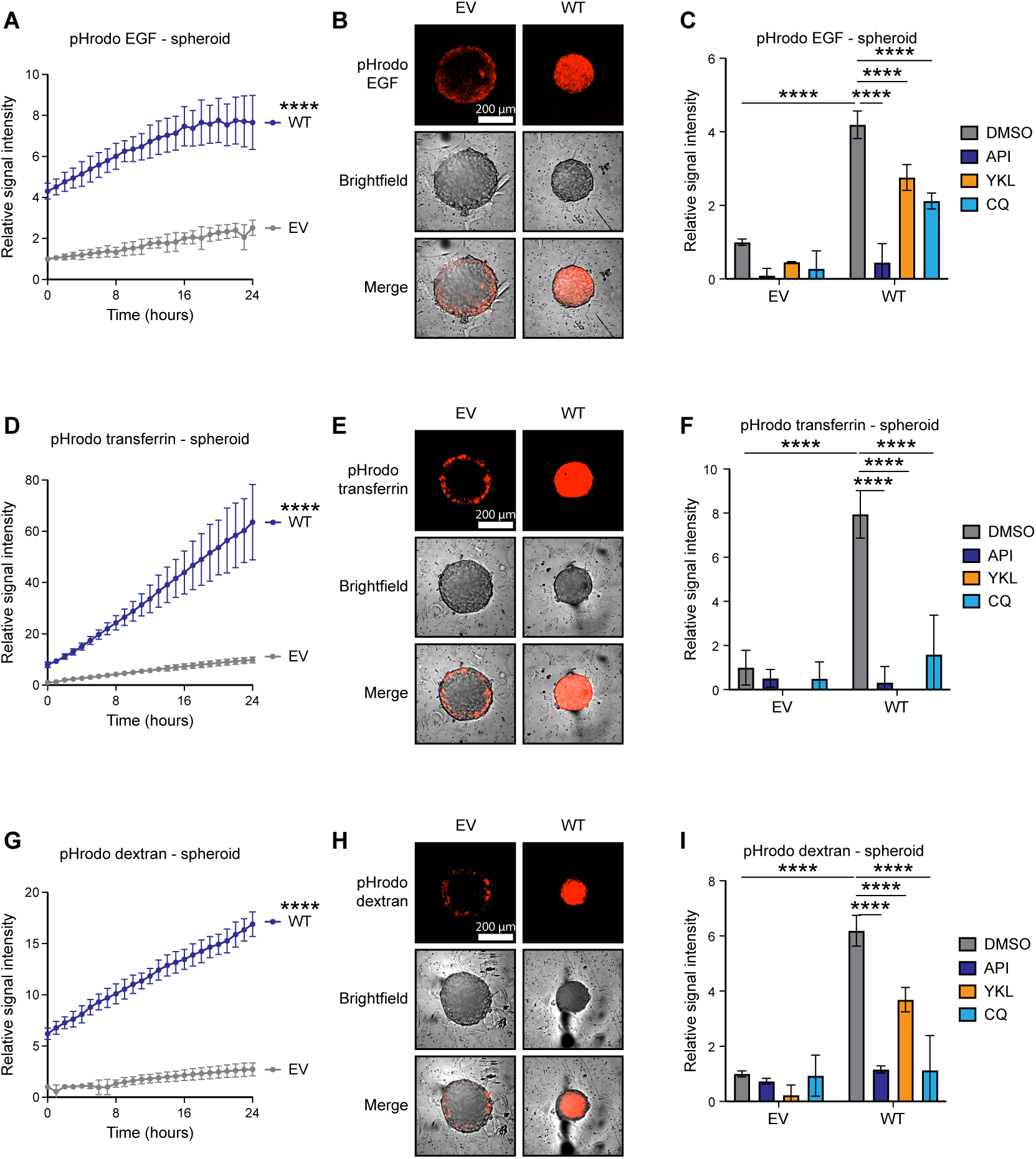
LKB1 promotes the endocytosis of cell surface receptors and extracellular dextran through PIKFYVE. **(A)** Relative signal intensity of A549 EV and WT spheroids treated with a pH-sensitive reporter that indicates the internalization of EGFR (pHrodo EGF). **(B)** Representative images of EV and WT spheroids treated with pHrodo EGF. **(C)** Relative signal intensity of EV and WT spheroids treated with DMSO, API 100 nM, YLK 2000 nM, or chloroquine (CQ) 10 µM and pHrodo EGF. **(D)** Relative signal intensity of A549 EV and WT spheroids treated with a pH-sensitive reporter of the internalization of transferrin (pHrodo transferrin). **(E)** Representative images of EV and WT spheroids treated with pHrodo transferrin. **(F)** Relative signal intensity of EV and WT spheroids treated with DMSO, API, YLK, or CQ and pHrodo transferrin. **(G)** Relative signal intensity of A549 EV and WT spheroids treated with a pH-sensitive reporter of the endocytosis of dextran (pHrodo dextran). **(H)** Representative images of EV and WT spheroids treated with pHrodo dextran. **(I)** Relative signal intensity of EV and WT spheroids treated with DMSO, API, YLK, or CQ and pHrodo dextran. Error bars indicate standard deviation. n = 3-4. **** indicates p<0.0001.

To ask whether LKB1 controls the internalization of EGFR or whether LKB1 influences the endocytosis of proteins and extracellular molecules more broadly, we evaluated the effect of LKB1 on the uptake of transferrin or dextran labeled with pHrodo. Similar to our findings with EGFR, we found that LKB1 promoted the endocytosis of transferrin and dextran. This effect was again reversed when the spheroids were pretreated with apilimod, with a SIK inhibitor, or with chloroquine (Figures 4D-I). Therefore, LKB1 promotes the endocytosis of cell surface receptors and extracellular polysaccharides in a SIK- and PIKFYVE-dependent manner.

### LKB1 impairs growth by opposing EGFR

We then sought to determine if LKB1-mediated growth suppression could be attributed to regulation of EGFR. To gauge whether this was likely, we first evaluated the rate of occurrence of mutations in *LKB1* and *EGFR*, since mutational patterns in tumor sequencing data can reveal pathways that cooperate or antagonize each other in tumorigenesis (Varmus et al., 2016). Upon our analysis of three sequencing studies of human LUAD tumors (Cerami et al., 2012), we found that mutations in *LKB1* and *EGFR* did not co-occur in any tumors (Figures 5A, B). This suggests that mutations in *LKB1* and *EGFR* are either functionally redundant or toxic when they co-occur, due to their regulatory effects on a common pathway.

**Figure 5:**
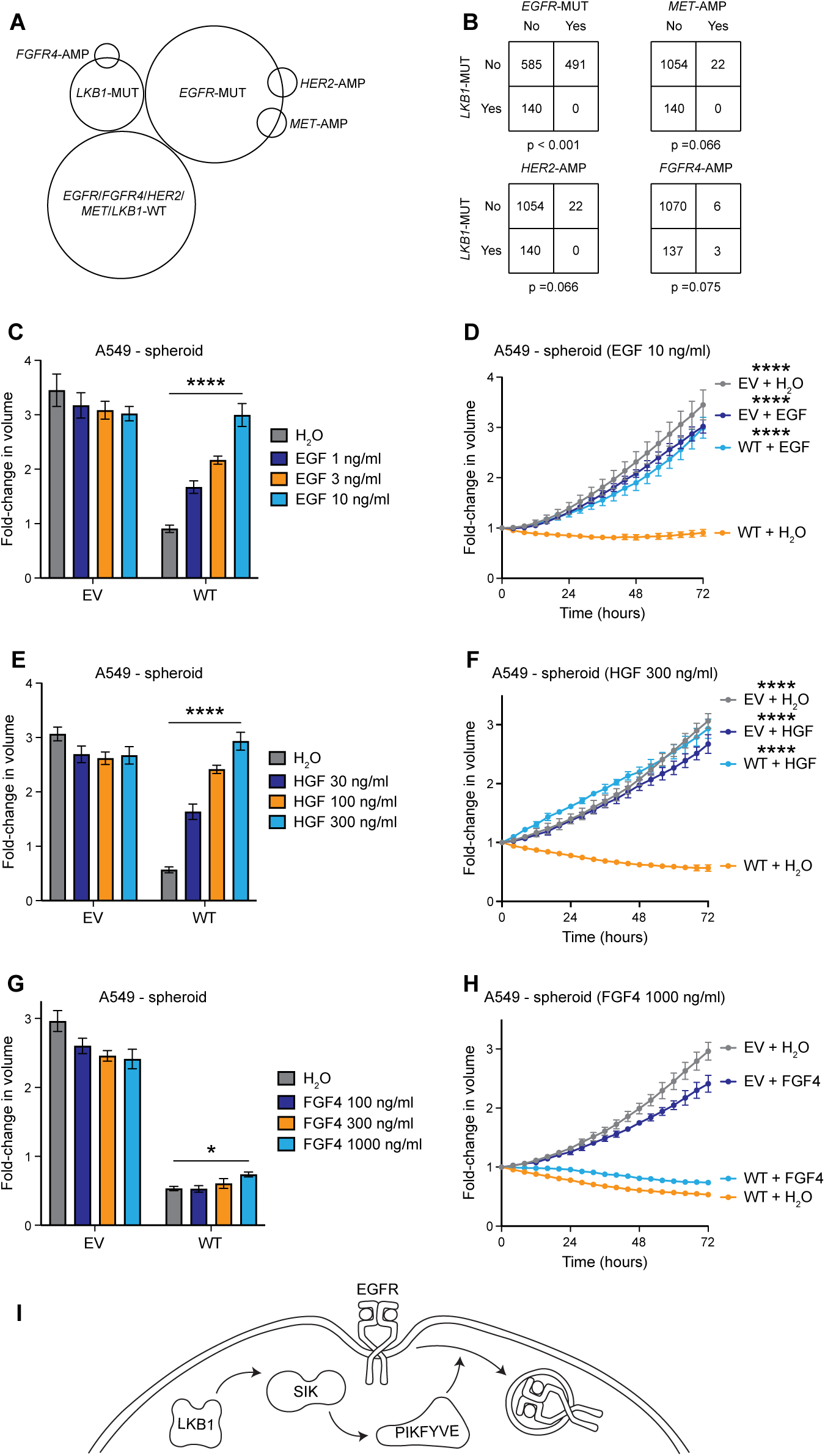
LKB1 suppresses growth by antagonizing the function of growth factor receptors. **(A, B)** Co-occurrences of mutations in *LKB1* and *EGFR* and amplifications of *HER2*, *FGFR4*, and *MET* in human LUAD tumors. **(C-H)** Growth of A549 EV and WT spheroids treated with H_2_O, EGF, HGF, or FGF4 at 72 hours in culture and in different doses of growth factors **(C, E, and G)**, or over a period of 72 hours in H_2_O or growth factors at the indicated doses **(D, F, and H)**. **(I)** A model of LKB1-mediated growth suppression. Error bars indicate standard deviation. n = 3-6. * and **** indicate p<0.05 and 0.0001. P-values indicate pairwise statistical comparisons to “WT + H_2_O” in panels **(D, F, and H)**.

Since a previous study showed that LKB1 influences the phosphorylation of not only EGFR, but also MET, HER2, and FGFR4 (Okon et al., 2014), we assessed whether alterations in these other receptors were mutually exclusive with mutations in *LKB1*. We found that amplifications in *MET* and *HER2* did not co-occur with mutations in *LKB1*, whereas amplifications in *FGFR4* did co-occur with mutations in *LKB1* (Figure 5A), but the frequencies at which these receptors were amplified were too small to determine whether the patterns were statistically significant (Figure 5B). Importantly, the addition of EGF or HGF to the culture media fully restored the growth of spheroids with WT *LKB1* in a dose-dependent manner, while having no effect on spheroids lacking LKB1 (Figures 5C-F). In contrast, even high doses of FGF4 had minimal effect on the growth of spheroids with WT *LKB1* (Figures 5G, H). Taken as a whole, these findings suggest that LKB1 inhibits growth by reducing the availability of WT EGFR and possibly other growth factor receptors, such as MET, on the cell surface. Thus, we propose that LKB1 inhibits pro-growth signaling by promoting the endocytosis of growth factor receptors via the SIKs and PIKFYVE (Figure 5I).

### Loss of *Lkb1* sensitizes *Kras*-mutant mouse lung tumor cells to combined inhibition of Egfr and Kras

Given that LKB1 may suppress growth by facilitating the internalization of EGFR, we wondered if tumors with mutations in *LKB1* may be driven by WT EGFR. Prior studies have shown that WT EGFR can augment the activity of oncogenic KRAS (Lito et al., 2016; Patricelli et al., 2016).

Oncogenic mutations in *KRAS* occur in approximately 50% of LUADs that harbor mutations in *LKB1* (Figures 6A, B). Therefore, in *KRAS*-mutant tumors, loss of LKB1 could promote the activity of KRAS through an increase in the activity of WT EGFR. Thus, we asked whether inhibition of EGFR would increase the sensitivity of *KRAS*-mutant lung cancer cells to inhibition of oncogenic KRAS. To do this, we used CRISPR/Cas9 to knock out *Lkb1* in a *Kras*^G12D^;*Trp53*^fl/fl^ mouse lung tumor line (634T, Figure 6C) (Liu et al., 2013b). We then treated these cells with various concentrations of the EGFR inhibitor erlotinib, along with the KRAS-G12D inhibitor MRTX1133 (Wang et al., 2022). We found that treatment of Lkb1-intact cells with erlotinib and MRTX1133 did not have a synergistic effect, as indicated by a drug synergy score (Bliss score = 3.32) (Zheng et al., 2022) that did not meet statistical significance (p = 0.235) (Figure 6D).

**Figure 6:**
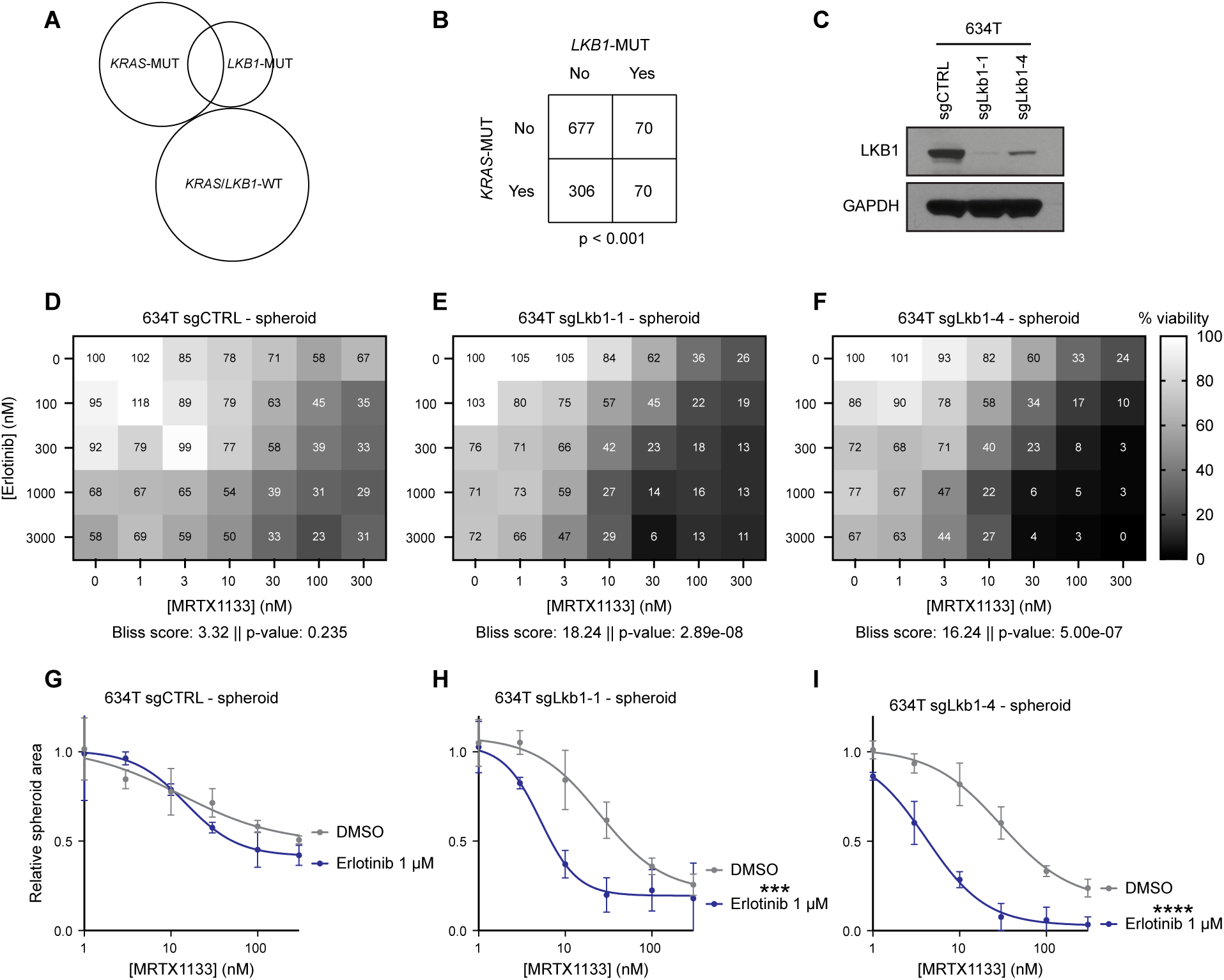
Loss of *Lkb1* sensitizes *Kras*-mutant murine lung tumor cells to combined inhibition of Egfr and Kras. **(A, B)** Co-occurrence of mutations in *LKB1* and *KRAS* in human LUAD tumors. **(C)** Lkb1 protein levels in a *Kras*^G12D^;*Trp53*^fl/fl^ mouse lung tumor line (634T) containing a control guide RNA (sgCTRL) or guide RNAs targeting *Lkb1*. **(D-F)** Dose-response matrices of 634T spheroids treated with erlotinib or the KRAS-G12D inhibitor MRTX1133 and containing sgCTRL **(D)** or sgRNAs against *Lkb1* **(E, F)**. The number listed over each condition corresponds to the percent viability of each spheroid relative to untreated spheroids (as measured by total spheroid area). Each condition represents the average percent viability of 3-4 spheroids. Additionally, drug synergy (Bliss) scores and their associated p-values are indicated below each matrix. **(G-I)** Dose-response curves of MRTX1133 in 634T spheroids containing sgCTRL **(G)** or sgRNAs against *Lkb1* **(H, I)**, in presence or absence of erlotinib 1 µM. Error bars indicate standard deviation. n = 3-4. *** and **** indicate p<0.0002 and 0.0001.

However, in cells containing a knockout of *Lkb1*, the combination of erlotinib and MRTX1133 had a clear synergistic effect, as indicated by Bliss scores that were statistically significant (Figure E, F). This observation was especially evident when viewing the effect of erlotinib 1 µM on the sensitivity of these cells to a range of doses of MRTX1133. Erlotinib had minimal effects on the sensitivity of Lkb1-intact cells to the Kras inhibitor (Figure 6G), but the efficacy of MRTX1133 was increased in cells with a knockout of *Lkb1* (Figures 6H, I). These results are consistent with the idea that loss of LKB1 drives the growth of lung tumor cells through an increase in the activity of KRAS mediated by WT EGFR.

## DISCUSSION

LKB1 controls a broad and incompletely defined signaling network, complicating efforts to identify the proteins responsible for the tumor suppressive activity of LKB1 – particularly those proteins positioned downstream of the SIKs. To address this challenge, we used an assay based in spheroid culture to study the tumor suppressive function of LKB1, which recapitulated the findings of prior studies conducted in animal models (Hollstein et al., 2019; Murray et al., 2019). This allowed us to perform an unbiased genome-wide CRISPR screen for genes required for growth suppression by LKB1. Additionally, the spheroid culture system enabled a detailed study of the function of LKB1 in human LUAD lines. The use of human cells is significant because LKB1-mediated signaling downstream of the SIKs is known to differ between mouse and human cells (Stein et al., 2023).

Our CRISPR screen identified two genes – *FIG4* and *VAC14*, both of which encode components of the PIKFYVE complex – as elements required for LKB1-mediated growth control. Moreover, chemical inhibition of PIKFYVE itself impaired growth suppression by LKB1. Since LKB1 is known to phosphorylate and activate PIKFYVE through AMPK – a kinase that is closely related to the SIKs – it is likely that LKB1-SIK may also activate the PIKFYVE complex through direct phosphorylation of PIKFYVE, as opposed to the other members of the complex, FIG4 and VAC14.

Additionally, we provide evidence that LKB1 – through PIKFYVE – may broadly activate the endocytosis of cell surface receptors and extracellular nutrients. This action of LKB1- PIKFYVE also results in the internalization of WT EGFR, which could impair growth by rendering a cell less responsive to EGF. Since LKB1 is generally thought to inhibit anabolic cellular processes when a cell is under nutrient stress (Shackelford and Shaw, 2009), a global upregulation of endocytosis by LKB1-PIKFYVE may serve this end by decreasing the responsiveness of a cell to growth factors, in order to limit cell growth and proliferation when nutrients are scarce.

We also note that PIKFYVE accounts for only part of the regulatory effect of LKB1 on EGFR. Prior studies have shown that LKB1 promotes the de-phosphorylation of growth factor receptors, including EGFR, by activating protein tyrosine phosphatases, such as PTPN12 (Okon et al., 2014). Therefore, LKB1 may inhibit WT EGFR through at least two distinct mechanisms: 1) by promoting PIKFYVE-dependent endocytosis of EGFR, and 2) by deactivating internalized EGFR via protein tyrosine phosphatases.

Lastly, our findings could provide an explanation for why mutations in *LKB1* and *KRAS* frequently co-occur. The loss of LKB1 may augment the activity of oncogenic KRAS through an increase in the activity of EGFR. Accordingly, this may render *KRAS*/*LKB1*-mutant cells more susceptible to combined inhibition of WT EGFR and oncogenic KRAS. Other studies have shown that inhibition of EGFR can potentiate the effects of inhibitors of oncogenic KRAS in cancer cells (Lito et al., 2016; Patricelli et al., 2016). Currently, two clinical trials are evaluating the combination of inhibition of WT EGFR and oncogenic KRAS in solid tumors (NCT04793958 and NCT05358249). Therefore, our findings suggest that close attention to the efficacy of this regimen in the subgroup of *LKB1*-mutant tumors is warranted.

## ACKNOWLEDGEMENTS

We thank Guoan Zhang (Department of Biochemistry, Weill Cornell Medicine) and the Proteomics & Metabolomics Core Facility at Weill Cornell Medicine for assistance with the proteomics experiments, Wei Li (Departments of Genomics and Precision Medicine and Pediatrics, George Washington University) for providing guidance on the analysis of the CRISPR screens, Kyuho Han (MEDIC Life Sciences) for providing guidance on culturing spheroids, and Long He (Department of Pharmacology, Weill Cornell Medicine) and Timothy McGraw (Department of Biochemistry, Weill Cornell Medicine) for critical discussions of the data. J.R.F. was supported by the Lee Cooperman Physician-Scientist Training Award of the Damon Runyon Cancer Research Foundation (DRG 18-18). E.E.G. was supported by the Kenneth G. and Elaine A. Langone Fellow Award of the Damon Runyon Cancer Research Foundation (DRG-2343-18). L.C.C. is supported by the National Cancer Institute (R35CA197588). N.E.S. is supported by the National Human Genome Research Institute (R01CA218668, R01CA279135, DP2HG010099, and R00HG008171). H.V. is the Lewis Thomas University Professor Fund, Weill Cornell Medicine.

## AUTHOR CONTRIBUTIONS

J.R.F. and H.V. conceived the study and wrote the manuscript. N.E.S. provided training and guidance in the design, execution, and analysis of the CRISPR screens. J.R.F., J.T., Y.Z., M.J.N., N.K., E.E.G., O.M., K.L., A.M., and M.M. performed experiments. All authors participated in the conception and design of experiments, interpretation of data, and review of the manuscript.

## METHODS

### Cell lines and culture media

All LUAD cell lines (A549, Calu-1, H1792, H2030, MOR, and SW1573) used in this study were obtained from ATCC or Sigma-Aldrich (MOR only). The 634T cells were a generous gift from the laboratory of Kwok Wong (NYU). Cells were cultured in RPMI-1640 (Corning 10-040-CV) supplemented with penicillin–streptomycin and 10% fetal bovine serum. Spheroids were grown in media supplemented with methylcellulose (Thermo Fisher Scientific M352-500). To prepare 100 ml of methylcellulose-containing media, 1.2 g of methylcellulose was autoclaved in a bottle containing a stir bar. One-hundred ml of media was warmed to 37 °C and then added to the methylcellulose. The methylcellulose media was then shaken vigorously and allowed to dissolve at 4 °C overnight, with stirring. Cells were maintained at 37 °C in a humidified incubator containing 5% CO2. All cell lines (and their derivatives) were confirmed to be mycoplasma-free using the MycoAlert mycoplasma detection kit (Lonza LT07- 218) at the beginning and upon completion of all experiments.

### Two-dimensional and spheroid growth assays

Two-dimensional (2D) growth assays were performed by seeding 5000 cells per well into flat-bottom 96-well plates (Corning 3603). For 2D growth assays in media with reduced nutrients or serum, 2500 cells per well were seeded into flat-bottom 96-well plates. Twenty-four hours later, the cells were rinsed twice with PBS and then medium containing variable levels of glucose, amino acids, or serum was added. Dialyzed FBS was used for all the experiments in which cells were grown with reduced levels of nutrients or serum. For 2D growth assays on plates with different levels of surface tension, 1.5 x 10^5^ cells were seeded into 6-well CytoSoft plates (Advanced Biomatrix 5190). Images of these cells were taken every 4 hours using an Incucyte Zoom or Incucyte S3 live cell imaging system.

Single spheroid growth assays were performed by seeding 100 cells per well (for H2030), 500 cells per well (for Calu-1, H1792, SW1573, and 634T cell lines), or 1000 cells per well (for A549 and MOR cell lines) into low-attachment, V-bottom 96-well plates (S-bio MS- 9096VZ) containing 100 µl of regular media per well. The plate was spun at 300 x g for 5 min and then placed in an incubator overnight. The following day, 80 µl of medium was slowly aspirated from each well, and then 80 µl of methylcellulose-containing medium was layered on top of each spheroid. For all 96-well assays, a total of 100 µl of medium (with or without added methylcellulose) was used per well. Images of live cells were taken every 4 hours using an Incucyte Zoom or Incucyte S3 live cell imaging system. The volume of an individual spheroid was derived from the total cross-sectional area of the spheroid. The start of each spheroid growth assay (T = 0 hours) was approximately 8 hours after the addition of methylcellulose. For all live cell imaging experiments, a minimum of 3 replicates were performed for each condition. Wells were excluded from analysis if the Incucyte failed to image a spheroid at any time point.

### Spheroid growth assays with inhibitors of EGFR and KRAS

Cells derived from the 634T line were seeded into low-attachment, V-bottom 96-well plates as described above. The following day, 80 µl of medium was slowly aspirated from each well, and then 80 µl of methylcellulose-containing medium was layered on top of each spheroid. Twenty-four hours after adding methylcellulose, erlotinib (MedChem Express HY-50896) and MRTX1133 (MedChem Express HY-134813) were diluted in RPMI and added to the spheroids. Plates were imaged in an Incucyte S3 live cell imaging system after 5 days of treatment. GFP signal was used to determine the area of a spheroid, and the area of the spheroid was used a surrogate for the number of viable cells within a spheroid. SynergyFinderPlus (https://tangsoftwarelab.shinyapps.io/synergyfinder/) was used to calculate Bliss synergy scores of erlotinib and MRTX1133 in the cell lines derived from 634T.

### Cloning and retrovirus preparation

Plasmids containing cDNAs encoding WT and KI *LKB1* alleles were synthesized by Twist Biosciences. The alleles also contain mutations that confer resistance to CRISPR/Cas9 editing (without changing the amino acid sequence) with all the sgRNAs against *LKB1* in the Toronto Knockout version 3 (TKOv3) CRISPR library (Addgene 125517, a gift from Jason Moffat). These alleles were then cloned into pBABE-GFP (Addgene 10668, a gift from William Hahn) or pBABE-Hygro (Addgene 1765, a gift from Hartmut Land) by first linearizing these backbones with EcoRI and then performing a Gibson assembly with NEBuilder HiFi DNA Assembly Master Mix (NEB E2621L). To generate the plasmid coding for PIKFYVE-N1939K, cDNA fragments for *PIKFYVE-N1939K* were synthesized by Twist Biosciences. These fragments were then cloned into pLV-EF1a-IRES-Puro (Addgene 85132, a gift from Tobias Meyer) by first linearizing the backbone with EcoRI and then performing a Gibson assembly. Plasmids coding for *PIK3CA-E545K* (Addgene 82881; a gift from Jesse Boehm, Matthew Meyerson, David Root) and *myrAKT1* (Addgene 64606, a gift from David Sabatini and Kris Wood) were purchased from Addgene, and the cDNAs encoding the *PIK3CA- E545K* and *myrAKT1* alleles were subcloned into pLV-EF1a-IRES-Puro via Gibson assembly.

Retrovirus vectors were generated by co-transfecting the *LKB1*-containing plasmids with the pCMV-VSV-G packaging plasmid (Addgene 8454, a gift from Robert Weinberg) into GP2-293 cells (Takara 631458). Lentivirus was generated by transfecting the *PIKFYVE*-containing plasmid with the packaging plasmids pMD2.G and psPAX2 (Addgene 12259 and 12260, gifts from Didier Trono) into 293FT cells (Invitrogen R70007) using Lipofectamine 3000 (Invitrogen L3000015). Media were changed 5h after transfection, and then the supernatant was collected 48h later.

### Editing with CRISPR/Cas9 and lentivirus production

To knock out a single gene, an sgRNA was cloned into the LentiCRISPRv2 plasmid (Addgene 52961, a gift from Feng Zhang) or the LentiCRISPRv2-GFP plasmid (Addgene 82416, a gift from David Feldser). To knockout a second gene in the same line, an sgRNA was cloned into the LRT2B plasmid (Addgene 110854, a gift from Lukas Dow). Lentivirus was generated from these plasmids using the procedure described above.

The sgRNA sequences used in this study are as follows: sgCTRL, GTCTGTATTTCAGTCTGTGA and GGTTGGATAAGGCTTAGAAA (only as a control for the knockout of a second gene in the same line); sgSIK1, ATGGTCGTGACAGTACTCCA; sgSIK3, GTGCTTGCAGATCTGCTCCA; sgFIG4-2, AACCGCTCGAAATAAGCCCG; sgFIG4-4, TGATGGGAGAGCCAAACCTC; sgVAC14-1, AAAGCGGAAGGTGGCAGCGC; sgVAC14-2, GCCCACCTTGCCCAGTGCGA; sgLkb1-1, CCAGGCCGTCAATCAGCTGG; and sgLkb1-4, GAACAATGCCCTGGCTGTGT.

### Inhibitors and growth factors

For all spheroid culture-based assays where inhibitors or growth factors were used, cells were seeded into medium containing the vehicle (DMSO or H_2_O), inhibitor, or growth factor. The following day, media were aspirated and replaced with methylcellulose-containing media that also contained the vehicle, inhibitor, or growth factor. The vehicle concentration was 0.1% for all experiments. Inhibitors used in these experiments are as follows: YKL-05-099 (MedChem Express HY-101147), apilimod (MedChem Express HY-14644), and chloroquine (MedChem Express HY-17589A). Growth factors used in these experiments are as follows: EGF (PeproTech AF-100-15), FGF4 (PeproTech AF-100-31), and HGF (PeproTech 100-39H).

### Mice and xenografts

Animal procedures were performed with the approval of the Weill Cornell Medicine IACUC. Tumor volume was not allowed to exceed 1000 mm^3^. Prior to implantation, cells were resuspended in PBS and mixed 1:1 with Matrigel (Corning 356231). For the A549 xenograft experiment, 10^6^ cells were injected into single flanks of 6-week-old, female athymic mice (Envigo). Caliper measurements were performed every 3 days to monitor tumor growth.

### Detection of total and phosphorylated proteins

For western blot experiments, protein lysates were prepared in RIPA buffer and quantified using a BCA protein assay (Thermo Scientific 23225). Proteins were separated on 4–12% NuPAGE Bis-Tris polyacrylamide gels (Invitrogen WG1402), transferred to a nitrocellulose membrane, and probed with antibodies against LKB1 (Santa Cruz sc-374334), GAPDH (CST 2118S), phospho-AMPKα T172 (CST 2535S), AMPKα (CST 5832S), phospho-ULK1 S555 (CST 5869S), ULK1 (CST 8054S), phospho-MEK1/2 S217/221 (CST 9154S), MEK1/2 (CST 8727S), phospho-ERK1/2 T202/Y204 (CST 4370S), ERK1/2 (CST 4695S), phospho-AKT T308 (CST 4056S), AKT1 (CST 2938S), phospho-S6 S240/244 (CST 5364S), S6 (CST 2217S), β-actin (CST 3700S), FIG4 (Novus NBP3-05130), and VAC14 (Sigma-Aldrich SAB4200074).

For the measurements of phospho-EGFR Y1068 and total EGFR in spheroids, an Alpha SureFire Ultra Multiplex Phospho-EGFR (Tyr1068) and Total EGFR Assay Kit (PerkinElmer MPSU-PTEGFR-K-HV) was used. To perform these measurements, spheroids were lysed 72h after the addition of the methylcellulose-containing media. For each condition, 48 spheroids were pooled, rinsed once with PBS, and then resuspended in 100 µl RIPA buffer. Spheroids were then sonicated and subjected to one freeze-thaw cycle prior to performing the assay. Six measurements were performed for each condition. The signal intensity for phospho-EGFR Y1068 for each well was normalized to the total EGFR signal intensity from the same well.

### CRISPR screens

Genome-wide screens were performed with the TKOv3 CRISPR library (Addgene 90294, a gift from Jason Moffat). Lentivirus was generated from the plasmid library using 293FT cells and Lipofectamine 3000-based transfection, as described above. For each line (EV and WT), approximately 30% of 144 x 10^6^ cells were infected with the TKOv3 library virus, to achieve an average 500-fold representation of the sgRNAs in each condition of the screen. Cells were then selected on puromycin for 5 days. Next, 35 x 10^6^ cells were seeded into either 2D or mass spheroid culture. For the 2D screen, cells were seeded into HYPERFlasks (Corning 10030). For the spheroid screen, cells were first resuspended in 36 ml of normal media, and then resuspended in 144 ml of media containing methylcellulose. The cell suspension was then distributed evenly over 245 mm^2^ square assay dishes (Corning 431111).

To passage the cells in mass spheroid culture, spheroids were spun at 800 x g for 10 min, washed with PBS, resuspended in Accutase (STEMCELL Technologies 07920), and allowed to incubate at room temperature for 10 min with gentle agitation. Cells were passaged every 3 days for 21 days, at which point genomic DNA was extracted from cells grown under each condition. sgRNA inserts were amplified with NEBNext High-Fidelity 2X PCR Master Mix (NEB M0541L). Samples were then pooled in equimolar concentrations, purified by gel electrophoresis, and sequenced with an Illumina HiSeq kit. Sequencing reads were trimmed and aligned to the TKOv3 library using Cutadapt and Bowtie. This alignment returned a table of raw reads, and the sgRNAs with less than 30 raw reads were excluded from further analysis. The raw reads were then analyzed with MAGeCK-MLE to obtain a beta-score for each gene (34).

The pathway enrichment analysis was performed with the pathfindR program in the R software package. sgRNAs with a beta score of >0.5 in the spheroids with WT *LKB1* and <0.5 in the *LKB1*-null spheroids were used as the input into pathfindR.

### Proteomics and kinase activity analyses

To analyze the proteome and phospho-proteome of *LKB1*-null and -WT cells, cells were seeded into 2D or spheroid culture, with 5 replicates per condition. After 48 hours in culture, protein lysates were prepared in RIPA buffer. Proteins were then precipitated, digested with trypsin, de-salted, and labeled with a TMTpro 16plex Label Reagent Set (Thermo Scientific A44520). Small aliquots of each TMT-labelled sample were mixed and analyzed by liquid chromatography-mass spectrometry (LC-MS) to evaluate peptide labelling and sample ratios. Based on those results, the remaining samples were mixed at equal ratios. The mixed, TMT-labelled samples were fractionated by RPLC into 12 fractions. Five percent of each fraction was used for expression profiling. The remaining 95% was enriched for phosphorylated peptides by using TiO2 beads. All samples were analyzed by LC-MS. The resulting data were searched against a protein database using MaxQuant software. These data were then processed and statistically analyzed using Perseus and R software. All samples were normalized to their median signal intensities to account for differences in sample loading. To account for changes in the total levels of proteins, the relative signal intensities of phospho- peptides in the phospho-proteomics datasets were subsequently normalized to the median signal intensities of the corresponding peptides in the total proteomics datasets. To identify phospho-peptides that varied in abundance between spheroid and 2D culture, the relative signal intensities in the spheroid phospho-proteomics datasets were normalized to the corresponding median signal intensities in the 2D phospho-proteomics datasets. The kinase activity analysis was performed by using the “Enrichment Analysis” option of the “Kinase Prediction” tool available at https://www.phosphosite.org/kinaseLibraryAction.

### Kinase prediction analyses

The kinase prediction analyses for serine and threonine residues on the PIKFYVE complex and PTPN12 were performed by using the “Score Site” option of the “Kinase Prediction” tool (see URL above). The data from these predictions were further analyzed and visualized in R.

### Measurement of endocytosis

Measurement of EGFR internalization was performed by using pHrodo Red EGF (Invitrogen P35374) at a concentration of 0.5 ug/ml, pHrodo Red transferrin (Invitrogen P35376) at 25 ug/ml, and pHrodo Red dextran (Invitrogen P10361) at 20 ug/ml. pHrodo-linked substrates were added to spheroids 24h after adding methylcellulose-containing media. The mean signal intensity for red fluorescence was measured with an Incucyte S3 and then normalized to the area of the spheroid.

### Analysis of human LUAD sequencing data

To assess the rates of occurrence of mutations in *LKB1* and *EGFR* and amplifications of *FGFR4*, *HER2*, and *MET*, the cBioPortal database was used to combine and analyze 3 studies of LUAD tumors (MSK, J Thoracic Oncology 2020; MSK, NPJ Precision Oncology 2021; MSK, 2021). Alterations of unknown significance were excluded from the analysis.

### Statistical analysis

Data were visualized, and statistical analyses were performed using GraphPad Prism 9 and R. P<0.05 was considered statistically significant. To evaluate for statistically significant differences amongst groups in the xenograft and spheroid growth assays, the area under the curve (AUC) was first determined for each condition. Next, a two-tailed, unpaired t-test was used to compare AUCs in experiments with two conditions. For experiments involving greater than two conditions, ANOVA was used to compare AUCs. Additionally, for experiments with greater than two conditions, the group corresponding to *LKB1*-WT (or *LKB1*- WT with sgCTRL, DMSO, or H_2_O) was used as the reference. The Dunnett method was used to correct for multiple hypothesis testing. To evaluate for statistically significant differences amongst conditions in the proteomics experiments, p-values were calculated using 2-sample t-test and the false discovery rate method was used to correct for multiple hypothesis testing. To evaluate for statistically significant differences amongst conditions in the LC3 degradation assays, the p-EGFR measurement assay, and the EGFR internalization assays, p-values were calculated by using a two-tailed, unpaired t-test for experiments with two conditions. For experiments involving greater than two conditions, ANOVA was used to determine the p-value. The Tukey method was used to correct for multiple hypothesis testing. For statistical analysis of the human LUAD sequencing data, p-values were calculated by using a two-tailed Fisher’s exact test.

---

**Figure S1:**
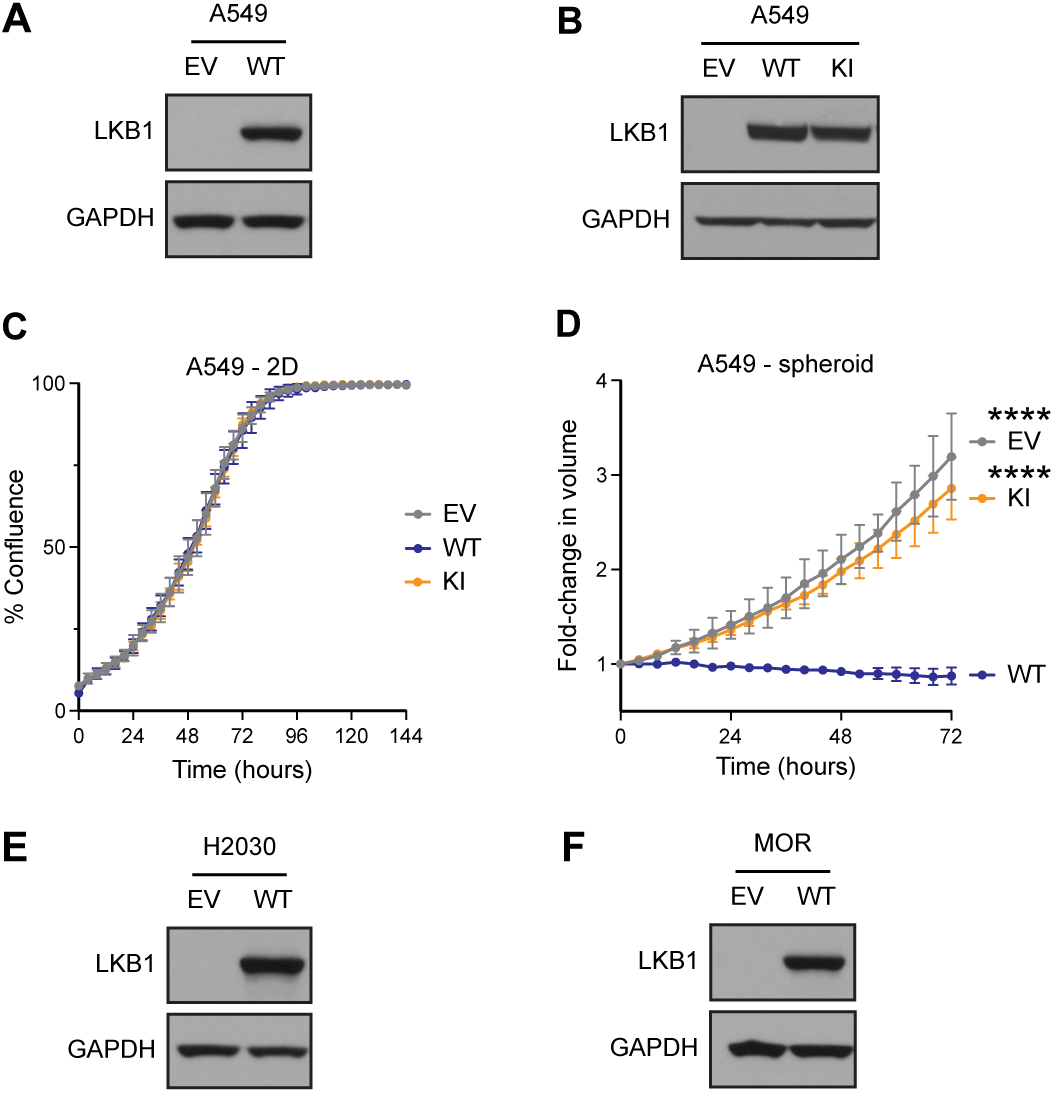
Expression of WT or kinase-inactive (KI) *LKB1* in *LKB1*-null LUAD lines. **(A, B)** LKB1 protein levels in EV, WT, and KI cells (derived from A549). **(C)** Growth of A549 EV, WT, and KI cells in 2D culture. **(D)** Growth of A549 EV, WT, and KI cells in spheroid culture. **(E, F)** LKB1 protein levels in EV and WT cells derived from H2030 and MOR LUAD lines.

**Figure S2:**
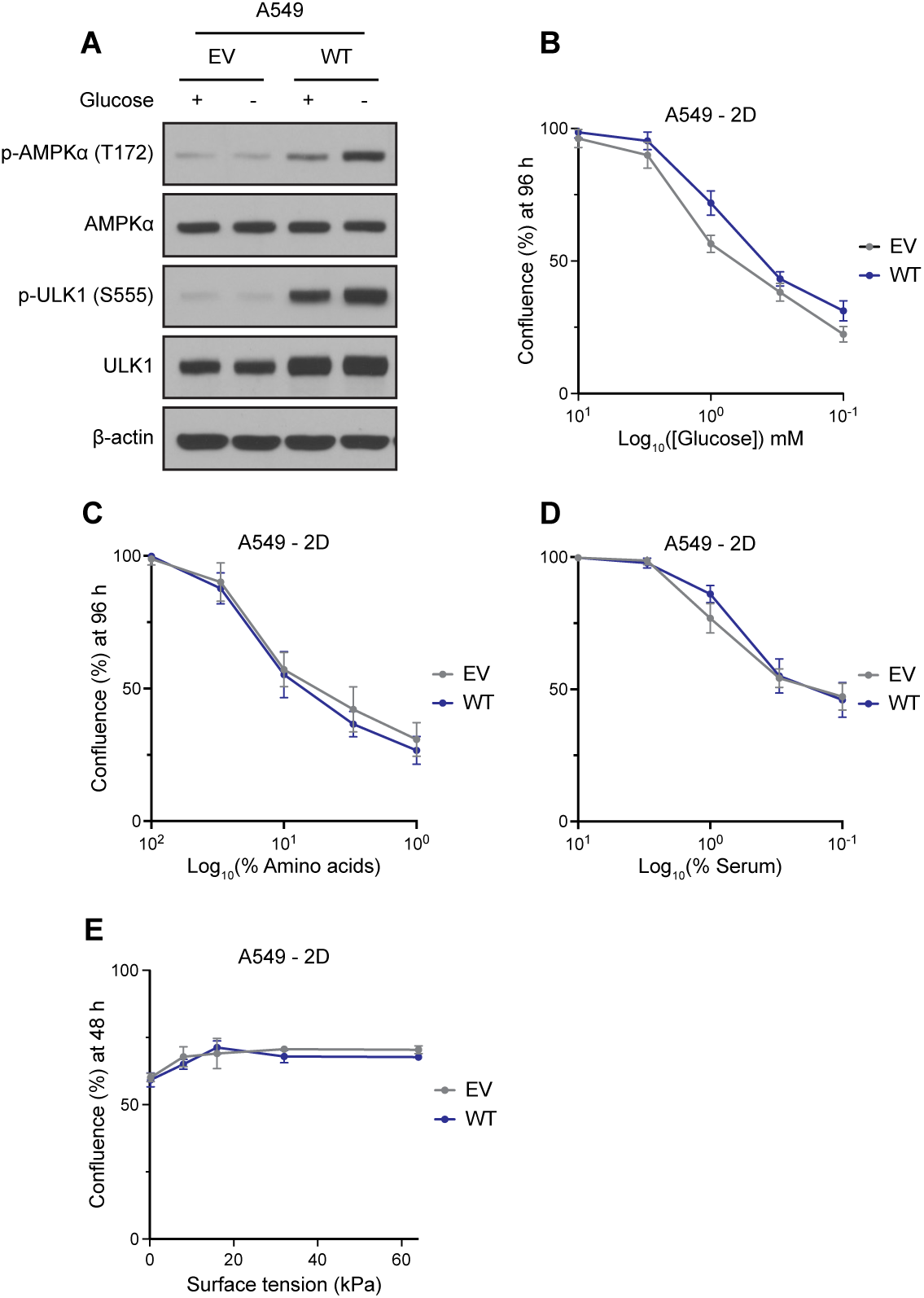
Nutrient deprivation, serum deprivation, or alterations in cellular surface tension do not recapitulate the tumor suppressive activity of LKB1 in 2D culture. **(A)** Levels of phosphorylated and total proteins in the LKB1-AMPK-ULK1 pathway from cells cultured in 2D, in media with or without glucose. **(B-E)** Growth of A549 EV and WT cells in 2D culture under variable levels of glucose **(B)**, amino acids **(C)**, serum **(D)**, or surface tension **(E)**. Error bars indicate standard deviation. n = 3-6.

**Figure S3:**
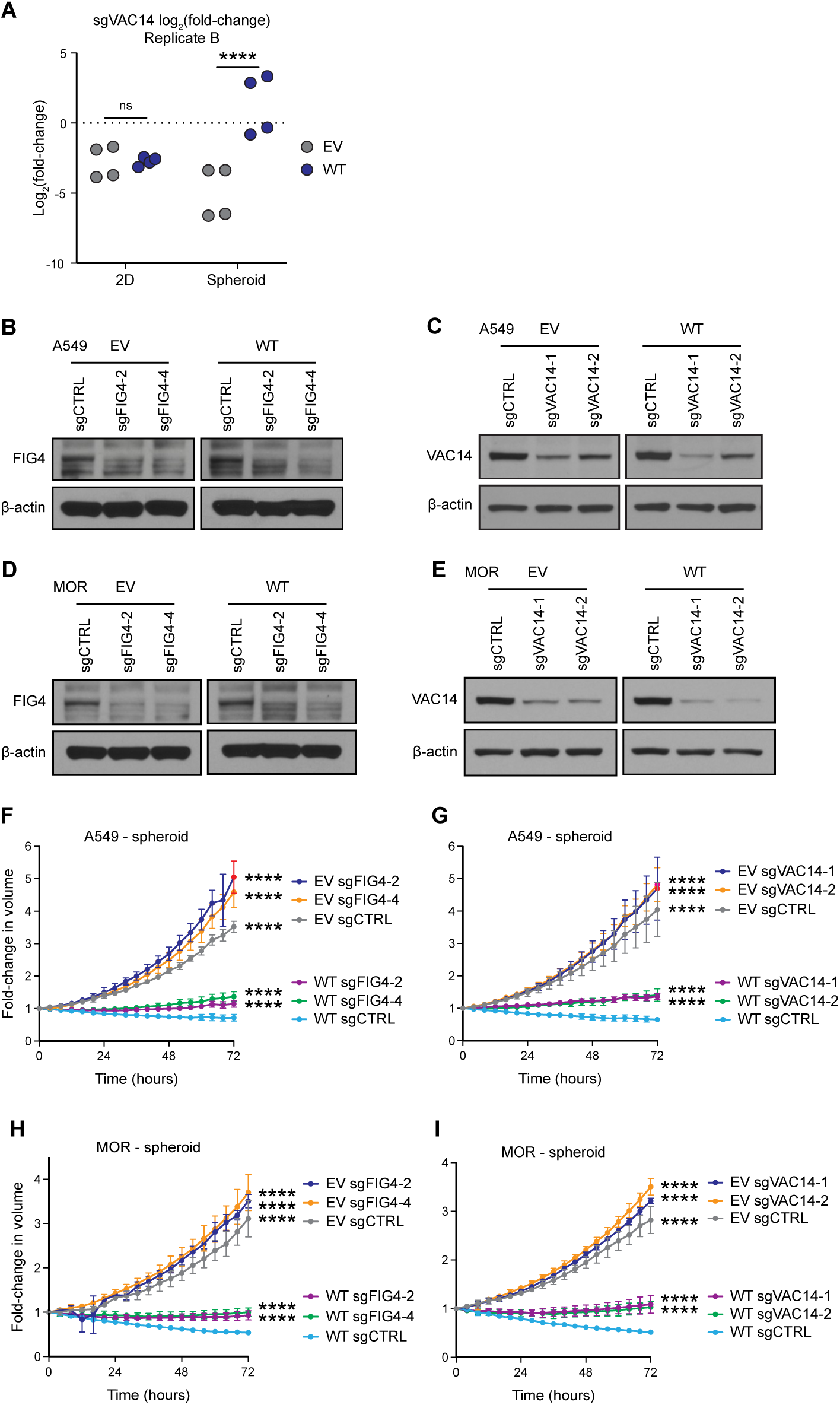
Validation of the genome-wide CRISPR screens. **(A)** Log_2_(fold-change) values of individual sgRNAs corresponding to *VAC14* in EV and WT cells – grown in 2D or spheroid culture – from a second replicate of the CRISPR screens. **(B-E)** FIG4 and VAC4 protein levels in EV and WT cells (derived from A549 or MOR lines) containing sgCTRL, or sgRNAs against *FIG4* or *VAC14*. **(F-I)** Spheroid growth of EV and WT cells (derived from A549 or MOR lines) containing sgCTRL, or sgRNAs against *FIG4* or *VAC14*. Error bars indicate standard deviation. n = 3-6. **** indicates p<0.0001. P-values indicate pairwise statistical comparisons to “WT sgCTRL.”

**Figure S4:**
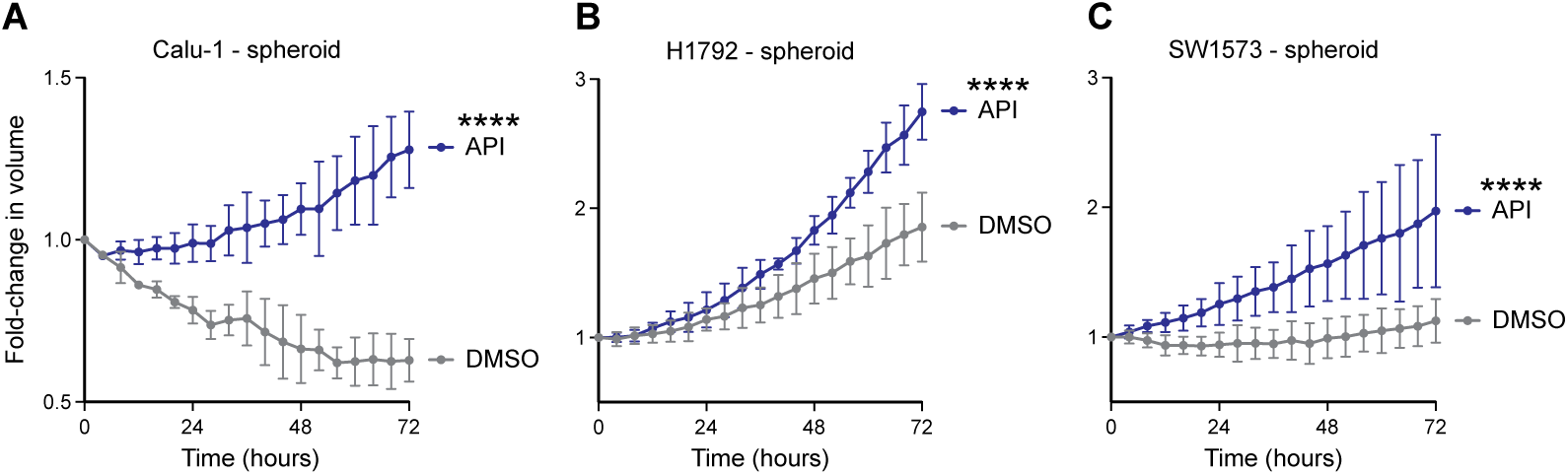
Inhibiting PIKFYVE promotes the growth of LUAD lines with endogenous WT *LKB1* as spheroids. **(A-C)** Spheroid growth of LUAD lines with endogenous WT *LKB1* (Calu-1 **(A)**, H1792 **(B)**, and SW1573 **(C)**) in the presence of DMSO or API. Error bars indicate standard deviation. n = 3-6. **** indicates p<0.0001.

**Figure S5:**
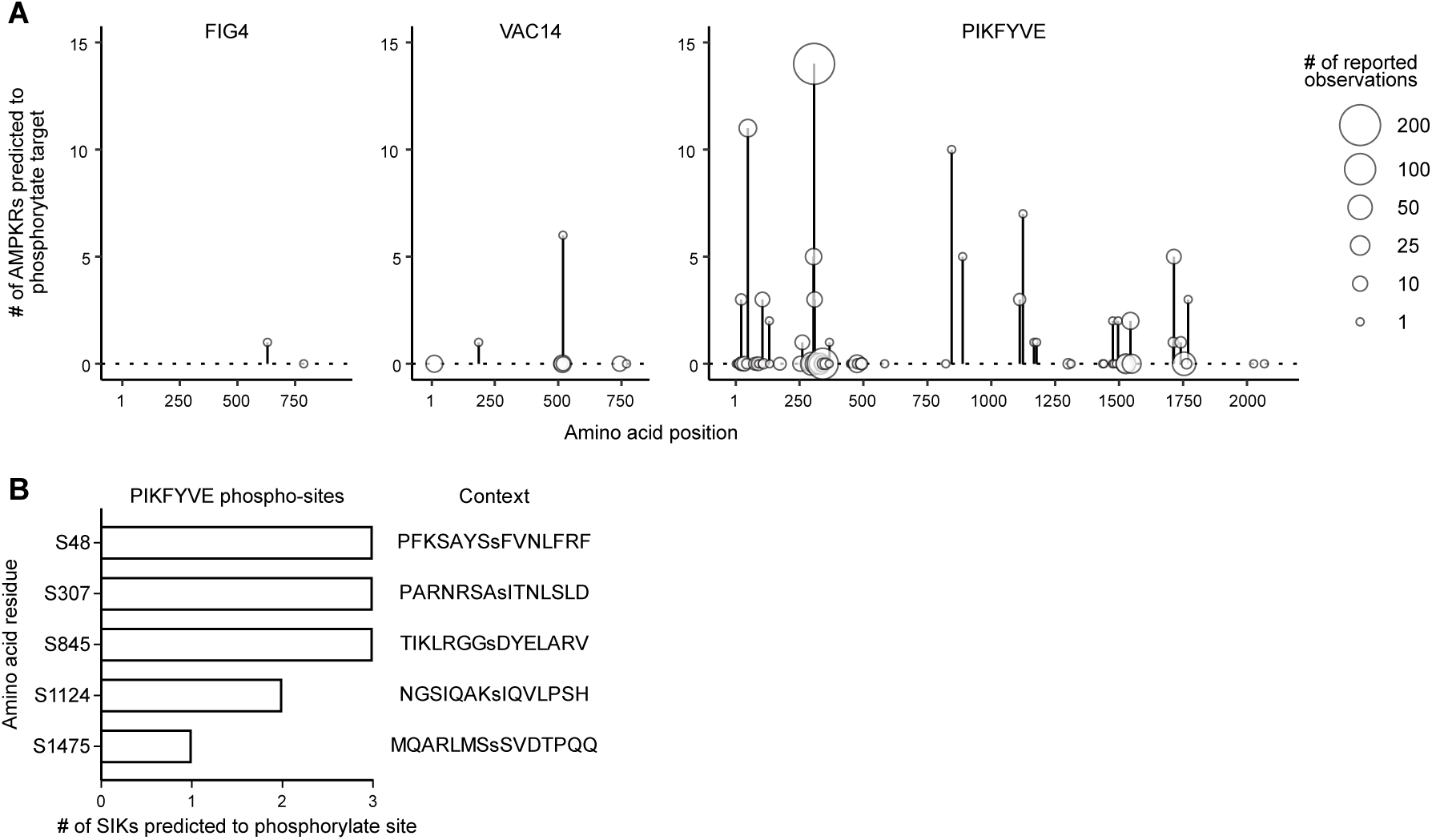
The SIKs phosphorylate PIKFYVE. **(A)** The number of AMPKRs predicted to phosphorylate serine and threonine residues within the PIKFYVE complex. The size of each circle indicates the number of times a phosphorylated site has been publicly reported. **(B)** Amino acid residues on PIKFYVE that are predicted to be phosphorylated by the SIKs.

## REFERENCES

Cai, X., Xu, Y., Cheung, A.K., Tomlinson, R.C., Alcazar-Roman, A., Murphy, L., Billich, A., Zhang, B., Feng, Y., Klumpp, M., et al. (2013). PIKfyve, a class III PI kinase, is the target of the small molecular IL-12/IL-23 inhibitor apilimod and a player in Toll-like receptor signaling. Chem Biol 20, 912–921.

Cerami, E., Gao, J., Dogrusoz, U., Gross, B.E., Sumer, S.O., Aksoy, B.A., Jacobsen, A., Byrne, C.J., Heuer, M.L., Larsson, E., et al. (2012). The cBio cancer genomics portal: an open platform for exploring multidimensional cancer genomics data. Cancer Discov 2, 401–404.

de Lartigue, J., Polson, H., Feldman, M., Shokat, K., Tooze, S.A., Urbe, S., and Clague, M.J. (2009). PIKfyve regulation of endosome-linked pathways. Traffic 10, 883–893.

Duex, J.E., Nau, J.J., Kauffman, E.J., and Weisman, L.S. (2006a). Phosphoinositide 5- phosphatase Fig 4p is required for both acute rise and subsequent fall in stress-induced phosphatidylinositol 3,5-bisphosphate levels. Eukaryot Cell 5, 723–731.

Duex, J.E., Tang, F., and Weisman, L.S. (2006b). The Vac14p-Fig4p complex acts independently of Vac7p and couples PI3,5P2 synthesis and turnover. J Cell Biol 172, 693–704.

Er, E.E., Mendoza, M.C., Mackey, A.M., Rameh, L.E., and Blenis, J. (2013). AKT facilitates EGFR trafficking and degradation by phosphorylating and activating PIKfyve. Sci Signal 6, ra45.

Gayle, S., Landrette, S., Beeharry, N., Conrad, C., Hernandez, M., Beckett, P., Ferguson, S.M., Mandelkern, T., Zheng, M., Xu, T., et al. (2017). Identification of apilimod as a first-in-class PIKfyve kinase inhibitor for treatment of B-cell non-Hodgkin lymphoma. Blood 129, 1768–1778.

Hart, T., Tong, A.H.Y., Chan, K., Van Leeuwen, J., Seetharaman, A., Aregger, M., Chandrashekhar, M., Hustedt, N., Seth, S., Noonan, A., et al. (2017). Evaluation and Design of Genome-Wide CRISPR/SpCas9 Knockout Screens. G3 (Bethesda) 7, 2719–2727.

Hollstein, P.E., Eichner, L.J., Brun, S.N., Kamireddy, A., Svensson, R.U., Vera, L.I., Ross, D.S., Rymoff, T.J., Hutchins, A., Galvez, H.M., et al. (2019). The AMPK-Related Kinases SIK1 and SIK3 Mediate Key Tumor-Suppressive Effects of LKB1 in NSCLC. Cancer Discov 9, 1606–1627.

Jin, N., Chow, C.Y., Liu, L., Zolov, S.N., Bronson, R., Davisson, M., Petersen, J.L., Zhang, Y., Park, S., Duex, J.E., et al. (2008). VAC14 nucleates a protein complex essential for the acute interconversion of PI3P and PI(3,5)P(2) in yeast and mouse. EMBO J 27, 3221–3234.

Johnson, J.L., Yaron, T.M., Huntsman, E.M., Kerelsky, A., Song, J., Regev, A., Lin, T.Y., Liberatore, K., Cizin, D.M., Cohen, B.M., et al. (2023). An atlas of substrate specificities for the human serine/threonine kinome. Nature 613, 759–766.

Krishnamurthy, N., Goodman, A.M., Barkauskas, D.A., and Kurzrock, R. (2021). STK11 alterations in the pan-cancer setting: prognostic and therapeutic implications. Eur J Cancer 148, 215–229.

Lees, J.A., Li, P., Kumar, N., Weisman, L.S., and Reinisch, K.M. (2020). Insights into Lysosomal PI(3,5)P2 Homeostasis from a Structural-Biochemical Analysis of the PIKfyve Lipid Kinase Complex. Mol Cell 80, 736–743 e734.

Lito, P., Solomon, M., Li, L.S., Hansen, R., and Rosen, N. (2016). Allele-specific inhibitors inactivate mutant KRAS G12C by a trapping mechanism. Science 351, 604–608.

Liu, Y., Lai, Y.C., Hill, E.V., Tyteca, D., Carpentier, S., Ingvaldsen, A., Vertommen, D., Lantier, L., Foretz, M., Dequiedt, F., et al. (2013a). Phosphatidylinositol 3-phosphate 5-kinase (PIKfyve) is an AMPK target participating in contraction-stimulated glucose uptake in skeletal muscle. Biochem J 455, 195–206.

Liu, Y., Marks, K., Cowley, G.S., Carretero, J., Liu, Q., Nieland, T.J., Xu, C., Cohoon, T.J., Gao, P., Zhang, Y., et al. (2013b). Metabolic and functional genomic studies identify deoxythymidylate kinase as a target in LKB1-mutant lung cancer. Cancer Discov 3, 870–879.

Mehenni, H., Gehrig, C., Nezu, J., Oku, A., Shimane, M., Rossier, C., Guex, N., Blouin, J.L., Scott, H.S., and Antonarakis, S.E. (1998). Loss of LKB1 kinase activity in Peutz-Jeghers syndrome, and evidence for allelic and locus heterogeneity. Am J Hum Genet 63, 1641–1650.

Mohseni, M., Sun, J., Lau, A., Curtis, S., Goldsmith, J., Fox, V.L., Wei, C., Frazier, M., Samson, O., Wong, K.K., et al. (2014). A genetic screen identifies an LKB1-MARK signalling axis controlling the Hippo-YAP pathway. Nat Cell Biol 16, 108–117.

Murray, C.W., Brady, J.J., Tsai, M.K., Li, C., Winters, I.P., Tang, R., Andrejka, L., Ma, R.K., Kunder, C.A., Chu, P., et al. (2019). An LKB1-SIK Axis Suppresses Lung Tumor Growth and Controls Differentiation. Cancer Discov 9, 1590–1605.

Okon, I.S., Coughlan, K.A., and Zou, M.H. (2014). Liver kinase B1 expression promotes phosphatase activity and abrogation of receptor tyrosine kinase phosphorylation in human cancer cells. J Biol Chem 289, 1639–1648.

Patricelli, M.P., Janes, M.R., Li, L.S., Hansen, R., Peters, U., Kessler, L.V., Chen, Y., Kucharski, J.M., Feng, J., Ely, T., et al. (2016). Selective Inhibition of Oncogenic KRAS Output with Small Molecules Targeting the Inactive State. Cancer Discov 6, 316–329.

Posor, Y., Jang, W., and Haucke, V. (2022). Phosphoinositides as membrane organizers. Nat Rev Mol Cell Biol 23, 797–816.

Shackelford, D.B., and Shaw, R.J. (2009). The LKB1-AMPK pathway: metabolism and growth control in tumour suppression. Nat Rev Cancer 9, 563–575.

Shisheva, A. (2001). PIKfyve: the road to PtdIns 5-P and PtdIns 3,5-P(2). Cell Biol Int 25, 1201–1206.

Stein, B.D., Ferrarone, J.R., Gardner, E.E., Chang, J.W., Wu, D., Hollstein, P.E., Liang, R.J., Yuan, M., Chen, Q., Coukos, J.S., et al. (2023). LKB1-Dependent Regulation of TPI1 Creates a Divergent Metabolic Liability between Human and Mouse Lung Adenocarcinoma. Cancer Discov 13, 1002–1025.

Sundberg, T.B., Liang, Y., Wu, H., Choi, H.G., Kim, N.D., Sim, T., Johannessen, L., Petrone, A., Khor, B., Graham, D.B., et al. (2016). Development of Chemical Probes for Investigation of Salt-Inducible Kinase Function in Vivo. ACS Chem Biol 11, 2105–2111.

Suprynowicz, F.A., Krawczyk, E., Hebert, J.D., Sudarshan, S.R., Simic, V., Kamonjoh, C.M., and Schlegel, R. (2010). The human papillomavirus type 16 E5 oncoprotein inhibits epidermal growth factor trafficking independently of endosome acidification. J Virol 84, 10619–10629.

Varmus, H., Unni, A.M., and Lockwood, W.W. (2016). How Cancer Genomics Drives Cancer Biology: Does Synthetic Lethality Explain Mutually Exclusive Oncogenic Mutations? Cold Spring Harb Symp Quant Biol 81, 247–255.

Vining, K.H., and Mooney, D.J. (2017). Mechanical forces direct stem cell behaviour in development and regeneration. Nat Rev Mol Cell Biol 18, 728–742.

Wang, B., Wang, M., Zhang, W., Xiao, T., Chen, C.H., Wu, A., Wu, F., Traugh, N., Wang, X., Li, Z., et al. (2019). Integrative analysis of pooled CRISPR genetic screens using MAGeCKFlute. Nat Protoc 14, 756–780.

Wang, X., Allen, S., Blake, J.F., Bowcut, V., Briere, D.M., Calinisan, A., Dahlke, J.R., Fell, J.B., Fischer, J.P., Gunn, R.J., et al. (2022). Identification of MRTX1133, a Noncovalent, Potent, and Selective KRAS(G12D) Inhibitor. J Med Chem 65, 3123–3133.

Zheng, S., Wang, W., Aldahdooh, J., Malyutina, A., Shadbahr, T., Tanoli, Z., Pessia, A., and Tang, J. (2022). SynergyFinder Plus: Toward Better Interpretation and Annotation of Drug Combination Screening Datasets. Genomics Proteomics Bioinformatics 20, 587–596.

